# Identification and calibration of ultrabright localizations to eliminate quantification error in SMLM

**DOI:** 10.1101/2022.04.06.487310

**Authors:** Bo Cao, Jielei Ni, Gang Niu, Danni Chen, Gang Liu, Lingxiao Zhou, Tingying Xia, Fu Feng, Shibiao Wei, Xiaocong Yuan, Yanxiang Ni

**Author notes:** **Correspondence: Yanxiang Ni**. These authors contributed equally: Bo Cao and Jielei Ni.

## Abstract

Single molecule localization microscopy (SMLM) is irreplaceable among super-resolution microscopies in revealing biological ultra-structures, given its unmatched high resolution. However, its sub-optimal quantitative capability, which is critical for characterizing true biomolecular organization of ultra-structures in cells, has hindered its widest application in biomedical research. Here, in SMLM imaging of cellular structures such as lipid rafts and microtubules with saturation labelling, we identified ultra-bright localizations, each of which is contributed by simultaneous emission of multiple molecules within a diffraction-limit region and has been regarded before as a regular localization from single molecule. Consistently, ultra-bright localizations are also observed in simulated SMLM imaging of endoplasmic reticulum or microtubules from public resource. Furthermore, after calibrating each ultrabright localization into multiple single-molecule localizations using the photon-number-based models, the density of total localizations shows linear correlation with the true molecule density, presenting SMLM with new reconstruction method as a quantitative analysis approach. Therefore, identification and dissection of ultra-bright localizations in SMLM enable the close and quantitative estimate of the true biomolecular organization.

## Introduction

Single molecule localization microscopy (SMLM) plays an irreplaceable role in characterizing biological ultra-structures, mainly due to its unmatched high resolutions of around 20 nm in the lateral dimension and 50 nm in the axial dimension^1-6^. In cells, distances of neighboring biomolecules range from one to a few nanometers, making it extremely challenging to display the true molecular organization of a certain ultrastructure. In this regard, SMLM has been intensively studied to technically approach biomolecular resolution^7-9^ and best image qualities^10-12^. Even though, SMLM meets great challenges in quantitative capability^13-19^, which hinders its widest application in biomedical research.

SMLM resolves close biomolecules within diffraction-limited region via stochastically activating their labeling photo-switchable fluorophores. To date, Alexa Fluor-647 (AF647) and Cy5 have been selected as the best dyes with high photon yields and low duty cycles (around 0.1%), each fluorophore switching between a long dark state and a short fluorescent state for various times during SMLM imaging. Ideally, point spread functions (PSFs) recorded in each camera frame for emitting fluorophores are sparse enough and separated from each other, so that each PSF is mostly contributed by and can be fit to determine the nanometer-precise localization of a single molecule. For those overlapping PSFs in densely labeling SMLM, localizations of molecules at a density of up to 8.8 emitters/*µ*m^2^ can be determined via different algorithms, such as compressed sensing and deep learning methods ^20-25^). In the final reconstructed SMLM image, the localizations of all molecules are commonly present to reflect the distribution and abundance of target biomolecules within the ultra-structures of interest, or to be segmented into cluster for molecular counting analysis. However, the switching times for each fluorophore during an image sequence stochastically vary from one to dozen, leading to a large uncertainty in localization number for a certain ultra-structure and probably accounting for sub-optimal quantitative capability of SMLM imaging.

In fact, cellular ultra-structures, like a vesicle with diameter of around 200 nm or a 200 nm-length segment of microtubule, for SMLM imaging usually contain dozens to hundreds of molecules. Such a large number of biomolecule number within a diffraction-limited region endows a relatively large probability of simultaneous emission of two or more molecules from this region. Given a 0.1% duty cycle, even this molecule number is only 50, there will be at least one special localization contributed by two or more close molecules within this region per one thousand frames. Such special localizations bring errors in localization quantitation has been regarded as most of them have been identified as regulator localizations contributed by single molecules. Therefore, we speculate the existence of special localizations contributed by multiple molecules and their possible effects on quantitative capability. This would be another factor that leads to the error of localization number compared to the truth. However, the calibration of such simultaneous emission of unresolvable multiple molecules hasn’t been exploit yet.

In this work, we demonstrated that such simultaneous emission of multiple close molecules, which we termed as ultra-bright localizations due to their high photon number, can widely exist in both 3D and 2D STORM imaging. The influence of ultra-bright localizations on quantitative analysis was evaluated, revealing the proportion of ultra-bright localizations can be xx with molecule density of xx. The probability density curves of photon number from ultra-bright localizations with different number of molecules was calculated. For an ultra-bright localization with certain photon number, the molecule number can be determined with the one with max probability. The calibrated molecule number is closely proportional to the true molecule number, providing access to quantitatively characterize molecule density. Statistical methods such as t-test were further introduced to overcome the error from uncertain blinking number of individual molecules.

## Results

### Ultra-bright localizations in 3D dSTORM imaging of densely labelled cellular samples

As the most-widely used dye for STORM, AF647 exhibits a duty cycle of as low as 0.1%, based on which AF647 has been generally regarded as an optional choice for STORM imaging of densely labelled subcellular structures. Given that sizes of most biomolecules are around a few nanometers, molecules within many sub-cellular structures in cells are relatively densely organized and profoundly abundant within a diffraction-limit region. For example, a single microtubule consisting of 13 protofilaments to form a 25 nm-wide hollow cylinder contains up to ∼ 375 *αβ*-dimers within an Abbe’s diffraction-limited image region, which is expected to exhibit a diameter of 231 nm with the use of an objective lens with a high numerical aperture of 1.4. Saturate labelling of alpha-tubulin or beta-tubulin will give more than 281 fluorescent molecules in a diffractionn-limited area, so that the probability of co-emission of more than one molecule for each frame will be larger than 3.27% considering a duty cycle of 0.1% and a survival fraction of 0.73. In this regard, it can be expected that, within 588 frames out of the STORM imaging sequence of 18000 frames, one co-emission from at least two fluorescent molecules can be detected on a segment of labeled microtubule in a single diffraction-limited area.

Co-emission of molecules within a diffraction-limited area poses great challenge to localization centroid determination algorithms. **Figure 1a** illustrates the emission spots and localization centroids of molecules separated by various distances. In STORM image reconstruction, centroids of emission spots completely separated from neighbouring spots are determined via fitting by a single 2D PSF (Gaussian function) (**I, Figure 1a**). Noises such as shot noise and pixelation noise will limit the precision of fitting, resulting in errors on the mean, width and ellipticity of the Gaussian function. When two molecules are closer but still farther than diffraction limit, their spots partially overlap but still resolvable (**II, Figure 1a**). Regular single-molecule fitting method would reject all such overlapping spots by different width and ellipticity filters. Recently developed high-density localization algorithms such as multi-molecule fitting may help localizing their centroids. However, when distance of two molecules is smaller than diffraction limit, the overlapped spots are unresolvable with a single peak (**III, Figure 1a**). Some of such spots probably can be recognized by multi-PSF fitting due to their large ellipticity. For some spots whose ellipticity and width fall within the range of ellipticity and width of single PSF, single-molecule fitting would fit the spot into a single 2D elliptical Gaussian function instead of rejecting it, leading to reduction of localization number not conforming to the true molecule number. For such simultaneous emission spots whose width and ellipticity are indistinguishable from those of single molecules, we defined them as ultra-bright spots, since most of them contain higher photon number and appear brighter than those contributed by single molecules.

**Figure 1.**
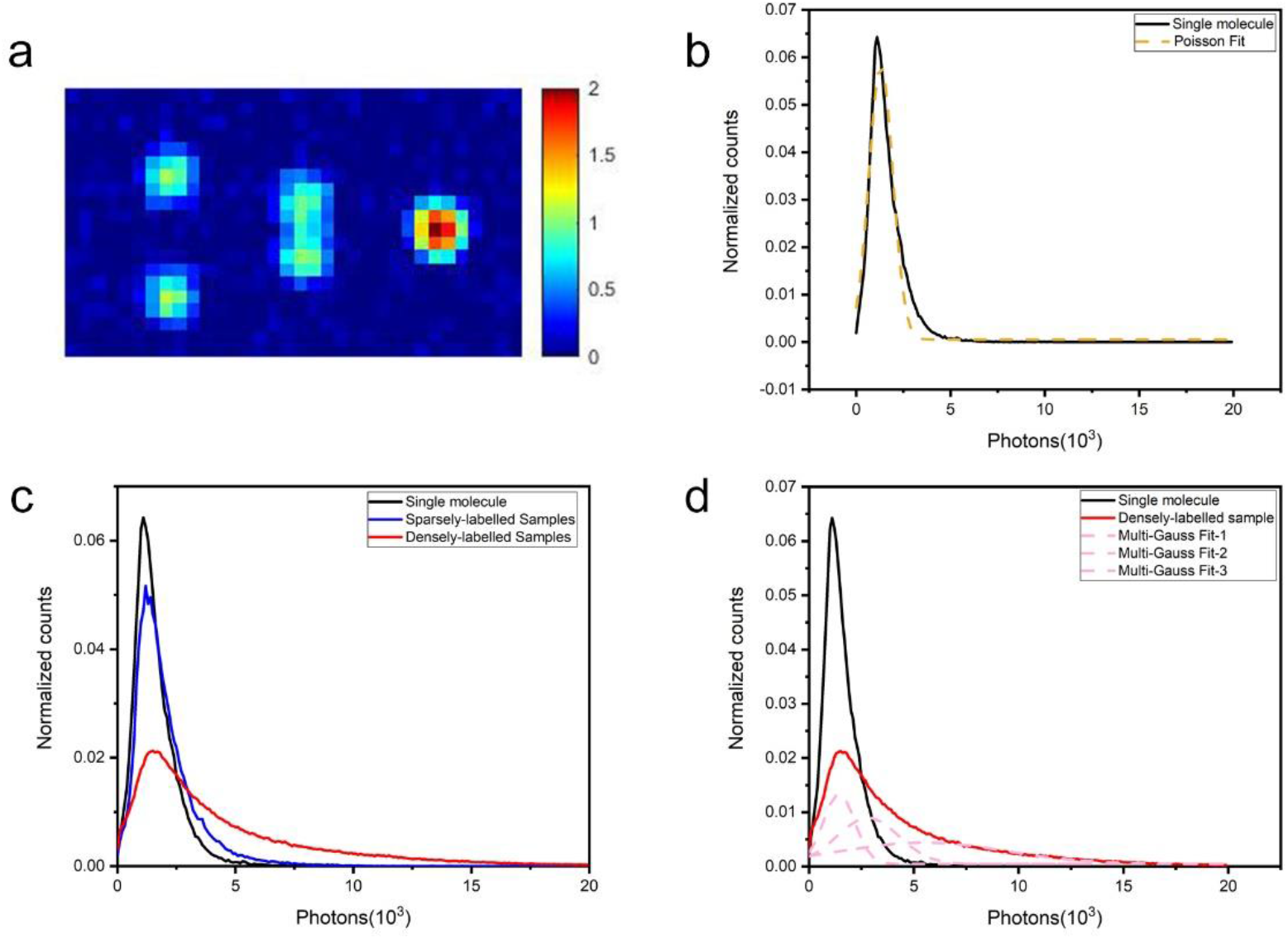
(a) The emission spots of two molecules with different distance. (I) Two molecules that are separated enough to one another produce separated spots, each of which can be localized by single-molecule fitting. (II) Two molecules that are close but hasn’t reached the diffraction limit produce overlapping spots that are resolvable. Such spots are usually rejected by single molecule fitting but can possibly be identified by algorithms such as multi-molecule fitting methods. (III) When two molecules are closer than diffraction limit, their spots are largely overlapped and unresolvable, which can be identified by single-molecule algorithm as a single molecule. (b) The probability density of photon number in sparsely distributing oligonucleotide sample, which can be fitted by a normal distribution. (c) The probability density curves of photon number from sparsely labeled (blue line) and densely labeled (red line) lipid raft show a distinct increase of high photon number portion with the increase of molecule density. The probability density of photon number from single oligonucleotides is also shown as the black line for a more intuitive comparison. (d) The probability density of photon number from densely labeled lipid raft can be fitted by a multi-Gaussian function (dashed red curve) with the left peak closely matches the peak from single oligonucleotides, indicating the existence of ultra-bright localizations.

To further verify the existence of ultra-bright spots in 3D dSTORM imaging of sub-cellular structures, we firstly characterized the photon number statistics of single molecules. Oligonucleotides each of which is conjugated to an AF647 molecule ^26-27^ were sparsely immobilized on glass surface to avoid overlapping of their emission spots. A total of 81839 localizations were collected after single-molecule fitting and width- and ellipticity-filtering to reject spots not matching the single PSF model well. After filtering, the width and ellipticity of these spots were shown in **Figure S1**. The black line in **Figure 1b** shows the probability density of photon number of single molecule spots calculated from the 81839 localizations. The probability density can be closely fitted by a normal distribution with a mean value of xx and standard deviation value of xx, as shown by the dashed yellow line in **Figure 1b**.

We then examined the difference of photon number of spots, which were all filtered by single-molecule algorithm, between sparsely labelling and densely labelling in plasma membrane lipid rafts. This ensures that both experiments, with densely-labelling and with sparsely-labelling, are under the same experimental condition except molecule density. Localizations were collected after single-molecule fitting and the same width- and ellipticity-filtering as above used. The width and ellipticity of the spots were shown in the bar graph in **Figure S1**, which shows no significant difference from those from single oligonucleotide molecules. The probability density of photon number in densely labelled lipid rafts is plotted as the red line in **Figure 1c**, in comparison of that from single oligonucleotide molecules which is also plotted here as the black line for a more intuitive comparison. As it is clearly seen from the two curves, the probability density of photon number in densely labelled lipid raft is distinctly characterized by a significant large portion of high photon number. In comparison, with the great decrease of molecule density, the probability density curve of sparsely labelled lipid raft, as the blue line in **Figure 1c**, approaches to that from single oligonucleotides (**Figure 1c**, black line). This supports that the high photon number portion in densely labelled lipid raft is contributed by the simultaneous emission of multiple close molecules.

In addition, we found that, the probability density of photon number from densely labelled lipid rafts can be fitted by a three-Gaussian function (**Figure 1d**, dashed red line, r square=0.998), with the left peak at similar value to that of single oligonucleotide molecules (**Figure 1d**, black line), supporting that other type of localizations is among all collected localizations.

### Ultra-bright localizations in 2D-STORM imaging

In 3D STORM with astigmatism imaging, the determination of the axial position of the molecule based on ellipticity of the spot can easily allow the generation ultra-bright localizations. For example, the co-emission of two close molecules in the focal plane, each of which is symmetric Gaussian spot, would overlaps into an elliptical spot and can be mistakenly identified as a molecule out-of-focus. Differently from 3D STORM, the point spread function in 2D STORM is usually more symmetric.

To further explore the existence of ultra-bright localizations in 2D STORM, we used the simulated endoplasmic reticulum datasets available on the EPFL SMLM challenge website (https://srm.epfl.ch/Challenge/ChallengeSimulatedData), which has been used for quantitative evaluation of different SMLM software ^28^. The dataset simulates endoplasmic reticulum structure in a field of view (FOV) of 6.4 × 6.4 × 0.7 μm^3^, with two different molecule densities (low molecule density of 0.2 per FOV and high molecule density of 5 per FOV). We used ThunderSTORM ^20^ to perform localization. After localization, each spot was extracted to fit with an elliptical Gaussian function. The fitted width 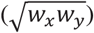, the ellipticity (*w*_*x*_/*w*_*y*_) and photon number of each spot were calculated. The barplot in **Figure 2a** compares the width (left pair of bars) and ellipticity (right pair of bars) of spots under two molecule densities, with the error bar representing standard deviation. The barplot shows that the width and ellipticity statistics of spots under high molecule density have no significant difference with those under low molecule density. However, as illustrating by **Figure 2b**, the probability density of photon number (calculated as the sum of grey values of the spot) under high density contains a distinct portion of high-photon-number spots (red line, **Figure 2c**), in comparison to that under low-density labelling (black line, **Figure 2c**), similar to our observation in 3D STORM. These results verify that ultra-bright localizations not only exist in 3D STORM but also can be generated in 2D STORM.

**Figure 2.**
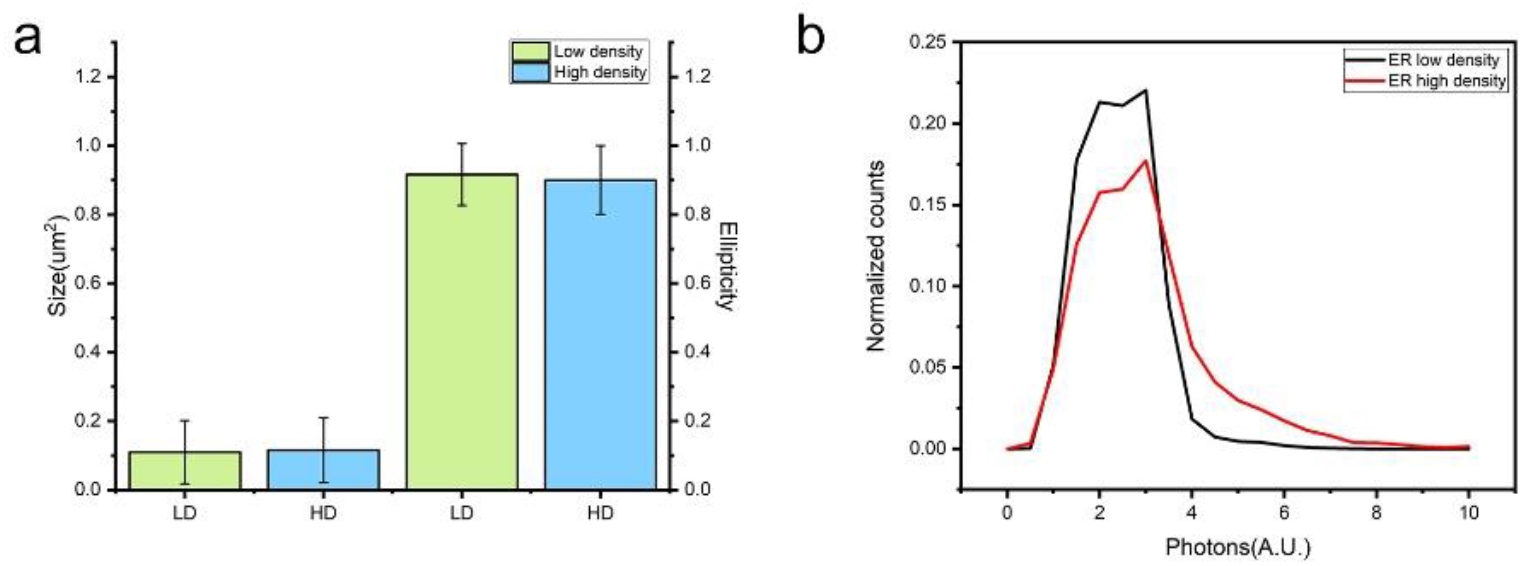
Comparison of width (a, left), ellipticity (a, right) and number of photons (b) of spots under low labeling density and high labeling density in simulated endoplasmic reticulum. Both width and ellipticity have no significant difference under two molecule densities (a). However, the probability density of photon number under high labeling density (red line) distinctly shows a larger portion of high photon number spots in comparison to that under low labeling density (black line) (b).

### Calibrating ultra-bright localizations allowing statistically quantitative analysis

Due to the wide existence of ultra-bright localizations in both 3D and 2D STORM imaging, localization number will be reduced severely with the increase of molecule density. To make a comparison of the number of ultrabright localization and true molecule number under different labelling density, we analysed a series of simulated images with different number of molecules randomly distributed in a diffraction region with a diameter of 231 nm. Molecule density ranges from 50 molecules to 450 molecules (5370 *µm*^−2^) in the simulation field. To ensure the simulation is close to real experiment, the blinking and photon statistics of each molecule are derived from our sparsely distributing oligonucleotides experiment (the dataset used in **Figure 1b**). Since the oligonucleotides are sparse enough, the emission spots of a molecule at different frame as well as the frame IDs of emission can be extracted from a series of 8000 frames. The position of the single molecule is determined by the centre of all the localizations it generated during 8000 frames. This serves as a single-molecule database for our simulation. Under a certain molecule density, we sampled randomly from the database with a certain number of molecules and putted them in the region with random positions. Then we performed localization of the 8000 frames using single molecule fitting algorithms and filtering as we used in **Figure 1b**. For each molecule density, the simulation was repeated 10 times to calculate the mean and standard deviation of localization number. We then compared the identified localization number to the true localization number, which is already known from the database. **Figure 3a** shows the true localization number increases linearly with molecule density, as the black line shows, while the identified localization number gradually deviates from the true localization number as the increase of molecule density, as the red line shows. The deviation originates from the formation of ultra-bright localizations within such diffraction limit region which cannot be resolved by algorithm.

**Figure 3.**
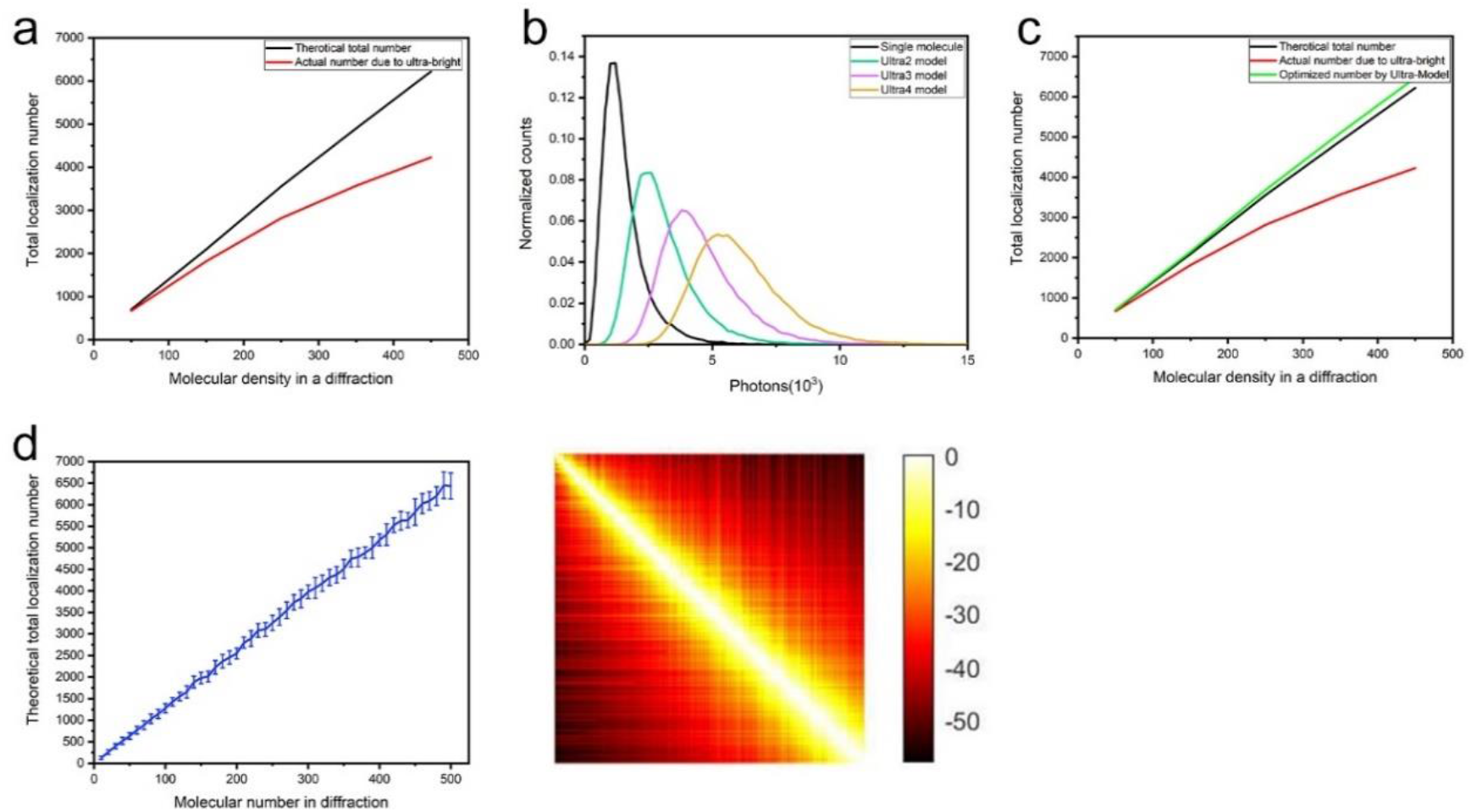
Calibration of ultrabright localization allows for quantitative analysis. (a) Comparison of the true localization number and the identified localization number localized by single emitter fitting within a simulated diffraction limit region with different molecule densities. The identified localization number decreases more as the increase of molecule density, due to the existence of ultra-bright localizations. (b) Probability density curves of photon number for different levels of ultra-bright localizations, in which Ultra-N stands for the simultaneous emission of N molecules. (c) the calibration of localization number of ultra-bright localizations, which is calculated as the molecule number of the level which has the maximum value of probability density, shows good linearity with molecule density and is close to the true localization numbers (green line). The localization number before calibration (red line) and the true localization number (black line) are also plotted for better comparison. (d) The standard deviations of the calibrated localization number with 20 simulations are shown as the error bars. (e) P-value map (log10-scale) of two-sample t-test of two different molecule densities, each of which range from 2-500 molecules within a diffraction limit region with a diameter of 231 nm.

To calibrate ultra-bright localization to its true localization number, we further investigated the photon number statistics of ultrabright spots generated by simultaneous emission of different numbers of molecules. We defined ultrabright spot that generated by N molecules as Ultra-N. For example, ultrabright spot that generated by two molecules is defined as Ultra-2. The green curve in **Figure 3b** shows the probability density of photon number of Ultra-2, which is simulated by randomly sampling from our single-molecule database with two spots. Similarly, we calculated the probability density of photon number of Ultra-3 and Ultra-4 as shown by the purple and yellow line in **Figure 3b**, respectively. The photon number distribution of single molecules is also plotted as black line in **Figure 3b** for better comparison. The probability density of higher level ultra-bright localization (Ultra-N with N>4) was not shown here. We can see from **Figure 3b** that the four probability density curves partially overlapped with each other. We then use these probability density curves to calibrate the ultra-bright localizations in **Figure 3a**. For each ultra-bright localization, we calculated the probability densities of different levels of ultrabright localizations (Ultra-N with N = 1, 2, 3, …) according to its photon number. The calibrated localization number was determined by the molecule number of the level which has the maximum value of probability density. The calibrated localization numbers at different molecule density are plotted in the green line in **Figure 3c**, which demonstrates good linearity with molecule density and is close to the line of true localization number (**Figure 3c**, black line).

The linearity of the calibrated localization number with molecule density provides access to quantitative analysis. For example, to determine the density change of target molecule under different treatment using STORM imaging, the change of molecule density usually reflected by localization number of the target molecule ^29^. Statistical techniques such as two-sample t-test can be used to determine whether two groups of density are equal. In **Figure 3d**, we plotted the p-value of two-sample t-test of pair-wise combination of molecule density. The simulated molecule number ranges from 2 to 500 within a diffraction limit region with a step of 2 molecules. For each molecule density, molecules are randomly selected from our single molecule database. Localization number under each molecule density is calibrated by the probability density model in **Figure 3c**. For each molecule density, the simulation was repeated 20 times. We then use two-sample t-test to calculate the p-value of two samples with sample size of 20. The p-value map in **Figure 3d** is shown in log10-scale. At low molecule density, even the difference of neighbouring density (the difference of density as low as 2 molecules in a diffraction limit region) can be statistically significant. At high molecule density, the minimum distinguishable density difference increases due to the increase of variation of localization number.

## Discussion

We have demonstrated that, ultra-bright localizations, which result from the co-emission of multiple molecules, commonly exist in STORM imaging of biological samples with sub-diffraction features, such as microtubule, lipid raft. The existence of ultra-bright localizations cannot be avoided due to the compromise of sufficient labelling that is required to sampling the fine structures and the limitation of dye whose duty cycle not low enough for dense labelling. Due to the existence of ultra-bright localizations, the localization density cannot linearly reflect the density of biomolecules, which becomes a crucial problem for quantitative analysis. We further demonstrated that, by modeling the photon number probability density of simultaneous emission of different numbers of molecules, ultra-bright localization can be successfully calibrated into the number of constituent molecules. The calibrated localization number is linearly increase with bio-molecule number. This provides a strong tool for quantitative analysis such as molecule abundance under different genetic and stoichiometry conditions, in which molecular densities under different condition or from a different region is usually compared.

Labelling with oligonucleotides is preferred in quantitative analysis, because it ensures only one fluorophore with one biomolecule. When using antibodies, a single protein can be labeled with multiple fluorophores, which increase the possibility of ultra-bright localization.

Although our results are demonstrated by dSTORM imaging, ultra-bright localizations are expected to exist in other SMLM approaches such as PALM, given their similar localization algorithms. In fact, PALM has been applied in a series of quantitative studies such as tracking the number of accessible phosphatidylinositol 3-phosphate binding sites in individual vesicles to reveal endosome maturation trajectory^18^, counting the centromere-specific histone H3 variant CENP-A^Cnp1^ in fission yeast for epigenetic inheritance studies ^30^. The calibration method of ultra-bright localizations that we proposed can also applied be in PALM experiments.

## Materials and methods

### dSTORM imaging system

The dSTORM system used in this study is based on an inverted optical microscope (IX-71, Olympus) with a 100× oil immersion objective lens (Olympus) as previously described [35]. A 641 nm laser (CUBE 640– 100C;Coherent) is used to excite fluorescence and switch AF647 to the dark state. The illumination uses the highly inclined and laminated optical sheet (HILO) configuration [36]. The laser power densities used this study is approximately 1.45 kW/cm^2^ for the 641 nm laser unless otherwise indicated. A dichroic mirror (ZT647rdc, Chroma) is used to separate the fluorescence from the laser and a band-pass filter (FF01-676/37, Semrock) on the imaging path is used to filter the fluorescence. Raw images of the fluorescent signals in each nuclear field are ac- quired with an EMCCD (DU-897U-CV0, Andor) at 33 Hz for 8000 frames. To avoid focal drift, an antidrift system is used to sustain the focal position within 10 nm during image processing [37].

### Sample preparation

To prepare single-molecule samples with immobilized oligonucleo- tides, cover glasses were cleaned by sonication for 15–25 min in water, washed with Mili-Q water, and then coated with 0.1% gelatin at room temperature for 10 min. Fluorescent microspheres (F8810, Thermo Fisher) of 200 nm in sizes were fixed on the gelatin-coated glasses with 4% paraformaldehyde for around 10 min at room temperature, so that they can act as fiducial markers to correct sample drift in the *x*–*y* plane during image acquisition. After being washed in PBS for multiple times, the glasses with beads were incubated for 30 min with 1 μM oligonucleotides, each of which was conjugated to an AF647 at its 5′ end and to biotin at its 3′ end, to allow for nonspecific association between biotin and gelatin, leading to immobilization of oligonucleotides sparsely on the glass surface. Subsequently, the samples were rinsed with PBS to remove unbound molecules, CTxB dilute in 1:30/1:300-900 respresented sparsely labeling and densely-labeling samples

### Cell sample preparation

The cells were detached with DMSO until the experiment. They were plated on pre-cleaned coverslips and cultured under DMSO for 24 hours. For CTB imaging, cells were washed three times with PBS and fixed with 4% PFA for 10 min. After washing out the fixing buffer with PBS three times, the sample was stained with 50 μl Alexa647-conjugated CTxb diluted in 1:30 or 1:900 and incubated in the dark for 20 min at RT. Finally the sample was washed with PBS three times.

### Image collection and processing

Samples on coverslips were embedded in dSTORM imaging buffer con- taining 50 mM Tris (pH 8.0), 10 mM NaCl, 1% β-mercaptoethanol (v/v), 10% glucose (w/v), 0.5 mg/mL glucose oxidase (G2133, Sigma), and 40 ug/mL catalase (C30, Sigma) [35, 38, 39]. Immediately after embed- ding, different samples of the same experiment set were subjected to dSTORM imaging field by field in turn to avoid any artificial difference caused by experiment condition changes with imaging time. For raw image analysis, a plugin Thunderstorm for Image J was applied.

## Data availability

The data that support the plots within this paper and other findings of this study are available from the corresponding authors upon reasonable request.

## Acknowledgements

This work was supported by Guangdong Major Project of Basic and Applied Basic Research (2020B0301030009), National Natural Science Foundation of China (31871293, 11774242, 62005180, 62105212), Shenzhen Science and Technology Planning Project (JCYJ20170817095211560), Shenzhen Peacock Plan (KQTD20170330110444030), Hefei National Laboratory for Physical Sciences at the Microscale (KF2020009). Natural Science Foundation of Guangdong Province (2016A030312010). X. Yuan appreciates the support given by the leading talents of Guangdong province (No. 00201505).

## Author Contributions

The manuscript was written through contributions of all authors. All authors have given approval to the final version of the manuscript.

## References

1. Betzig, E. et al. Imaging Intracellular Fluorescent Proteins at Nanometer Resolution. Science 313, 1642–1645 (2006).

2. Hess, S., Girirajan, T. & Mason, M. Ultra-High Resolution Imaging by Fluorescence Photoactivation Localization Microscopy. Biophysical Journal 91, 4258–4272 (2006).

3. Rust, M., Bates, M. & Zhuang, X. Sub-diffraction-limit imaging by stochastic optical reconstruction microscopy (STORM). Nature Methods 3, 793–796 (2006).

4. Heilemann, M. et al. Subdiffraction-Resolution Fluorescence Imaging with Conventional Fluorescent Probes. Angewandte Chemie International Edition 47, 6172–6176 (2008).

5. Schermelleh, L. et al. Super-resolution microscopy demystified. Nature Cell Biology 21, 72–84 (2019).

6. Lelek, M. et al. Single-molecule localization microscopy. Nature Reviews Methods Primers 1, (2021).

7. Gu, L. et al. Molecular resolution imaging by repetitive optical selective exposure. Nature Methods 16, 1114–1118 (2019).

8. Cnossen, J. et al. Localization microscopy at doubled precision with patterned illumination. Nature Methods 17, 59–63 (2019).

9. Balzarotti, F. et al. Nanometer resolution imaging and tracking of fluorescent molecules with minimal photon fluxes. Science 355, 606–612 (2016).

10. Schnitzbauer, J., Strauss, M., Schlichthaerle, T., Schueder, F. & Jungmann, R. Super-resolution microscopy with DNA-PAINT. Nature Protocols 12, 1198–1228 (2017).

11. Diekmann, R. et al. Optimizing imaging speed and excitation intensity for single-molecule localization microscopy. Nature Methods 17, 909–912 (2020).

12. Strauss, S. et al. Modified aptamers enable quantitative sub-10-nm cellular DNA-PAINT imaging. Nature Methods 15, 685–688 (2018).

13. Huijben, T. et al. Detecting structural heterogeneity in single-molecule localization microscopy data. Nature Communications 12, (2021).

14. Khater, I., Nabi, I. & Hamarneh, G. A Review of Super-Resolution Single-Molecule Localization Microscopy Cluster Analysis and Quantification Methods. Patterns 1, 100038 (2020).

15. Nicovich, P., Owen, D. & Gaus, K. Turning single-molecule localization microscopy into a quantitative bioanalytical tool. Nature Protocols 12, 453–460 (2017).

16. Shivanandan, A., Deschout, H., Scarselli, M. & Radenovic, A. Challenges in quantitative single molecule localization microscopy. FEBS Letters 588, 3595–3602 (2014).

17. Lee, S., Shin, J., Lee, A. & Bustamante, C. Counting single photoactivatable fluorescent molecules by photoactivated localization microscopy (PALM). Proceedings of the National Academy of Sciences 109, 17436–17441 (2012).

18. Puchner, E., Walter, J., Kasper, R., Huang, B. & Lim, W. Counting molecules in single organelles with superresolution microscopy allows tracking of the endosome maturation trajectory. Proceedings of the National Academy of Sciences 110, 16015–16020 (2013).

19. Rollins, G., Shin, J., Bustamante, C. & Pressé, S. Stochastic approach to the molecular counting problem in superresolution microscopy. Proceedings of the National Academy of Sciences 112, (2014).

20. Ovesný, M., Křížek, P., Borkovec, J., Švindrych, Z. & Hagen, G. ThunderSTORM: a comprehensive ImageJ plug-in for PALM and STORM data analysis and super-resolution imaging. Bioinformatics 30, 2389–2390 (2014).

21. Holden, S., Uphoff, S. & Kapanidis, A. DAOSTORM: an algorithm for high-density superresolution microscopy. Nature Methods 8, 279–280 (2011).

22. Nehme, E., Weiss, L., Michaeli, T. & Shechtman, Y. Deep-STORM: super-resolution single-molecule microscopy by deep learning. Optica 5, 458 (2018).

23. Nehme, E. et al. DeepSTORM3D: dense 3D localization microscopy and PSF design by deep learning. Nature Methods 17, 734–740 (2020).

24. Zhu, L., Zhang, W., Elnatan, D. & Huang, B. Faster STORM using compressed sensing. Nature Methods 9, 721–723 (2012).

25. Marsh, R. et al. Artifact-free high-density localization microscopy analysis. Nature Methods 15, 689–692 (2018).

26. Ni, Y. et al. Super-resolution imaging of a 2.5 kb non-repetitive DNA in situ in the nuclear genome using molecular beacon probes. eLife 6, e21660 (2017).

27. Ni, J. et al. Improved localization precision via restricting confined biomolecule stochastic motion in single-molecule localization microscopy. Nanophotonics 11, 53–65 (2021).

28. Sage, D. et al. Quantitative evaluation of software packages for single-molecule localization microscopy. Nature Methods 12, 717–724 (2015).

29. Zhang, H. et al. Reversible phase separation of HSF1 is required for an acute transcriptional response during heat shock. Nature Cell Biology 24, 340–352 (2022).

30. Lando, D. et al. Quantitative single-molecule microscopy reveals that CENP-A^Cnp1^ deposition occurs during G2 in fission yeast. Open Biology 2, 120078 (2012).

